# Projections of non-invasive human recordings into state space show unfolding of spontaneous and over-trained choice

**DOI:** 10.1101/2020.02.24.962290

**Authors:** Yu Takagi, Laurence Hunt, Mark W Woolrich, Timothy EJ Behrens, Miriam C Klein-Flügge

## Abstract

Choices rely on a transformation of sensory inputs into motor responses. Using invasive single neuron recordings, the evolution of a choice process has been tracked by projecting population neural responses into state spaces. Here we develop an approach that allows us to recover state space trajectories on a millisecond timescale in non-invasive human recordings. We selectively suppress activity related to relevant and irrelevant sensory inputs and response direction in magnetoencephalography data acquired during context-dependent choices. Recordings from premotor cortex show a smooth progression from sensory input encoding to response encoding. In contrast to previous macaque recordings, information related to choice-irrelevant features is represented more weakly than choice-relevant sensory information. To test whether this mechanistic difference between species is caused by extensive overtraining common in non-human primate studies, we trained humans on >20,000 trials of the task. Choice-irrelevant features were still weaker than relevant features in premotor cortex after overtraining.

## Introduction

For many decades, neuroscientists have studied task-dependent response properties of individual neurons and this has laid the groundwork for our current understanding of brain function. More recently, a major shift from looking at individual neurons to studying population responses has begun to shed light on larger-scale neural dynamics, thus providing insight into previously hidden circuit mechanisms ^1, 2^. Usually in this approach, the major axes of variation of a neural population are defined and the population activity is projected into this neural state space. This new way of looking at neural firing rates has begun to revolutionize our understanding of various neural processes, including the evolution of choice in parietal and frontal cortices ^3–6^, the critical stages of movement preparation and reaching in premotor and motor cortices ^7–9^, and the mechanisms underlying working memory in prefrontal cortex (PFC, ^10^).

Yet deriving neural population trajectories requires invasive recordings because the population vector is constructed from the firing rate of individual neurons. As a consequence, to date, no comparable non-invasive techniques for use in healthy human participants have been established. Here we asked whether we can recover neural population trajectories in humans, and thus provide insight into the unfolding of choice on a millisecond basis, by selectively suppressing neural populations to different features using repetition suppression in magnetoencephalography (MEG) recordings.

Repetition suppression (or “adaptation”) takes advantage of a feature first observed over 50 years ago ^11^, that the activity of a neuron will be suppressed when it is repeatedly exposed to features it is sensitive to. This phenomenon has since been widely replicated and shown to be a robust property of neurons in single-unit recordings (for review see ^12^). Importantly, the bulk signal measured from thousands or millions of neurons using techniques such as EEG, MEG or fMRI will also show suppression if a subset of these neurons is sensitive to a repeated feature. Thus, repetition suppression provides insight into the activity of specific subpopulations of neurons in these non-invasive recordings available for use in humans, and in that way allows us to examine the underlying neural mechanisms. Here, we extend the repetition suppression framework in one crucial way: we suppress the MEG signal to multiple different features within the same experiment. Each feature can be seen as one axis in a multi-dimensional state space and thus we ask whether repetition suppression along multiple features can mimic projections onto multiple axes in state space. If so, this would enable us to observe the equivalent of state space trajectories in humans over time, with temporal resolution in the order of milliseconds thanks to the temporal precision of MEG.

Our first key result shows that it is possible, using repetition suppression, to record neural population trajectories in humans. The recovered human population traces are similar to those obtained from invasive recordings in non-human primates (NHPs) ^4^, showing a progression from an encoding of choice inputs to an encoding of the motor response.

Once the feasibility of our non-invasive population approach was established, we asked whether the mechanisms of choice uncovered in NHPs also generalize to humans. More precisely, we examined whether the mechanisms for input-selection during choice were comparable between the two species. A large body of evidence shows that selection of relevant sensory inputs occurs through top-down modulations from prefrontal and parietal regions onto early sensory regions ^13–27^. However, two recent perceptual decision making studies in macaques found that irrelevant sensory inputs are not filtered out before the integration stage ^4, 28^. We speculated that top-down influences might be more important when someone is naïve at doing the task (see also ^29^). Untrained humans thus performed context-dependent choices in the same perceptual decision-making task used in macaques. This showed that information about both relevant and irrelevant input dimensions was present in premotor cortex, but irrelevant inputs were weaker than relevant inputs, consistent with top-down suppression.

One major difference in terms of how human and animal studies are conducted, however, is that animals are trained for thousands of trials before neural recordings are performed. Differences in input selection mechanisms could therefore be a consequence of overtraining. In an attempt to reconcile our findings with those obtained in macaques, we mirrored NHP training conditions and our human participants underwent extensive training on over 20,000 trials of the task. However, this did not change the input selection mechanisms evident in the MEG recordings taken afterwards. Irrelevant inputs were present, but still more weakly than relevant inputs. This was true when examining information encoded in premotor cortex, or when decoding from whole-brain activity.

## Results

Participants (n=22) performed a modified version of a dynamic random-dot-motion (RDM) decision task, in which stimuli simultaneously contained information about colour (red vs green) and motion (left vs right, as in ^4^). An additional flanker stimulus displayed 150ms before the RDM stimulus instructed participants about the relevant stimulus dimension: arrows indicated the choice in the current trial was about the dominant motion direction, whereas coloured dots instructed participants to respond based on the dominant colour (**Fig 1A**).

**Figure 1.**
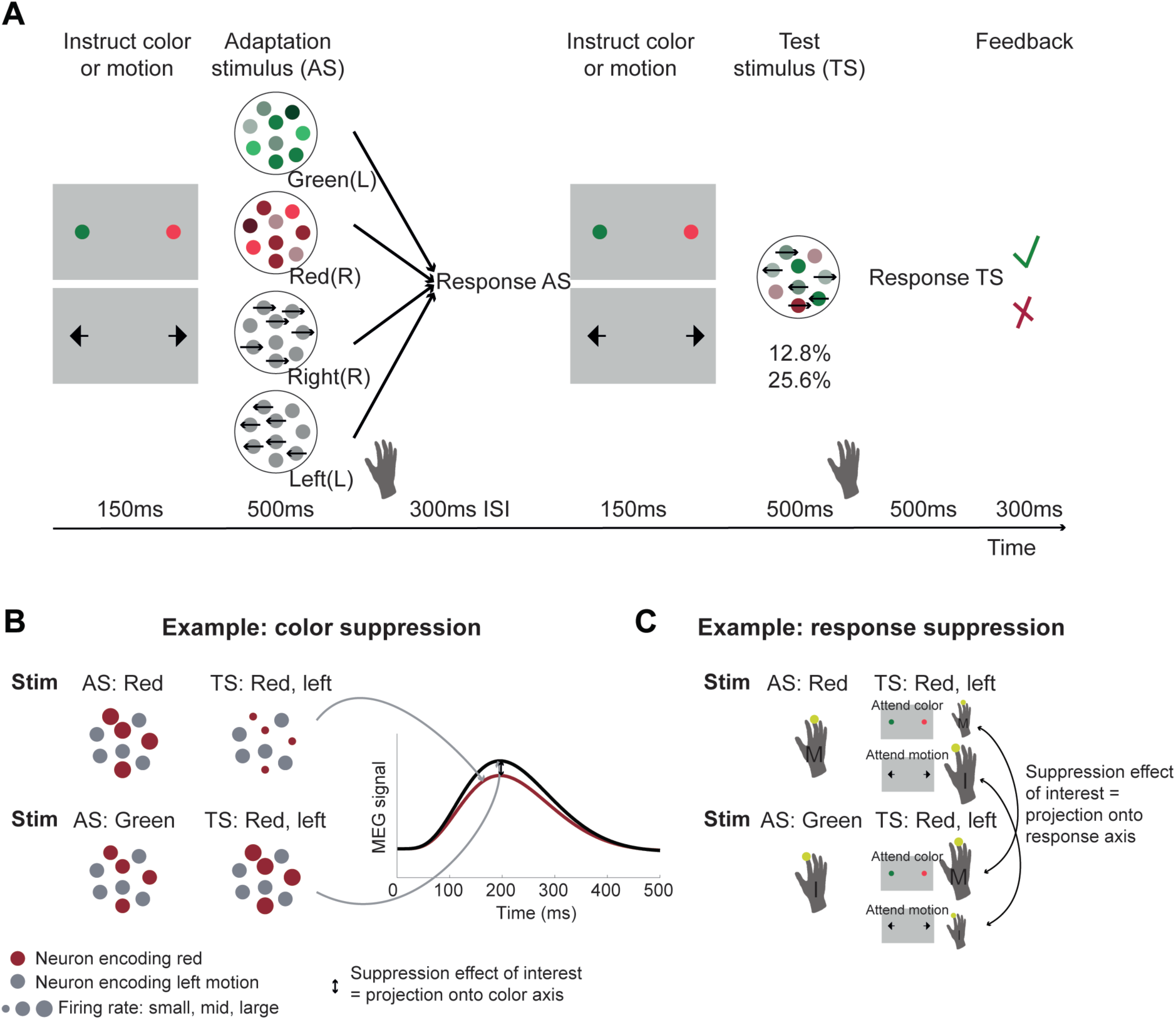
Experimental task involving manipulation of specific choice-features. **A**, Human participants performed a perceptual choice task adapted from the macaque version in ^77^ while MEG data was recorded. Each trial involved two random-dot-motion stimuli – an adaptation stimulus (AS) and a test stimulus (TS). A flanker cue (coloured dots or arrows) instructed which choice dimension, colour or motion, to attend to for making a choice. Responses were given using the right-hand index or middle finger for green/left and red/right stimuli, respectively. By presenting two choices with varying features in rapid succession, we selectively suppressed the subset of neurons sensitive to repeated features. To maximise suppression effects, AS colour and motion was strong compared to the TS (70% compared to 25.6% or 12.8% motion coherence or colour dominance). Feedback at the end of each trial related to performance on the TS. In total there were 64 conditions: 4 AS x 2 contexts x 2 directions x 2 colors x 2 coherence levels. The rationale for the selective suppression of choice features is illustrated for two examples in B and C. **B**, The top and bottom row show two combinations of AS and TS that were compared to extract colour suppression. At the time of the TS (the focus of all analyses), the stimulus is identical, containing predominantly red colour and left-ward motion. If preceded by a red AS (top row), any red-coding neurons will show a reduced signal at the time of the TS (red dots), but any other neurons will show the same response (grey dots). Thus, the overall MEG signal will be reduced compared to a situation where the preceding AS stimulus does not share any features with the TS (green; bottom row). This suppression effect, i.e. the difference in the MEG signal for two identical TS as a function of their preceding AS (arrow) can be captured in a time-resolved manner, thus showing not only whether but also *when* colour is encoded. This mimics a projection of the neural population onto the axis capturing variation in the encoding of colour, as commonly done in invasive neural recordings. **C**, The sequence of stimuli shown in B can also be used to probe response suppression. When participants are attending to colour at the time of the TS, a middle finger response will be repeated in the top example but not in the bottom. If they are attending to motion, an index finger response will be repeated in the bottom but not the top example. The respective differences (arrows) thus provide a time-resolved measure of response suppression, equivalent to projecting neurons onto a response axis.

### MEG activity can track the temporal evolution of a choice process

When recording activity from premotor cortex in non-human primates (NHPs) performing the same task, neural activity shows a clear evolution from an initial processing of the choice input (e.g. colour in a colour trial) to a later processing of the choice direction (motor response: right or left; ^4^). Our first analysis therefore aimed to establish whether the time-resolved nature of magnetoencephalography (MEG) data would allow us, in a similar way, to watch how a choice evolved from the processing of inputs to the processing of the motor response in human premotor cortex.

In order to get a handle on neurons that process colour, motion or response in MEG recordings measuring the summed activity of many neurons, we incorporated an additional task manipulation. The main RDM stimulus on each trial (‘test stimulus’: TS) was preceded by another RDM stimulus, the ‘adapting stimulus’ (AS), so that each trial contained two RDM stimuli in quick succession (**Fig 1A**). Unlike the TS, the AS contained only one input feature (strong red or strong green colour, or strong leftward or strong rightward motion). Participants responded to both stimuli using the corresponding finger of the right hand (index finger for leftward motion or green colour, middle finger for rightward motion or red colour). We hypothesised that presentation of the AS meant that MEG activity recorded at the time of the TS would have suppressed encoding for features already engaged at the time of the AS. For example, when a red and leftward TS was preceded by a red AS, MEG activity measured at the time of the TS would still contain the activity of neurons responding to leftward motion, but neurons responding to the red colour input would be suppressed, compared to a situation where the AS was green (**Fig 1B**). Furthermore, if participants were asked to attend to the colour of the TS, they would produce a middle finger response twice in quick succession (to AS and TS, both red), and thus the ‘response-direction (middle finger)’ selective neurons would also be suppressed (**Fig 1C**). This would not be the case for the identical stimuli when leftward motion was attended at the time of the test stimulus. In this case, only colour-sensitive neurons but not response-selective neurons would be selectively suppressed at the time of the TS. In summary, by creating and comparing situations with and without suppression for input (colour or motion) and response, we aimed to establish whether we could measure premotor cortex activity transition from an encoding of input to an encoding of choice output using non-invasive human MEG.

We focussed this analysis on the time-course of activity from an area in left premotor cortex (PMd) which was identified using a beamforming contrast capturing suppression to any input (colour or motion) or response at any timepoint between [-250,750]ms around the test stimulus (see Online Methods). This contrast was thus agnostic to any differences between input and response suppression, and agnostic to any timing differences between these effects. PMd, the premotor region responsible for selecting hand motor responses, was our *a priori* region of interest for this study. Its role in selecting finger responses is equivalent to the role of the frontal eye fields for guiding eye movement responses, the region where NHP recordings were performed ^4^. Indeed, left PMd, in a cluster together with left M1, was the strongest peak at the whole-brain level for this contrast (p(FWE)=.018 cluster-level corrected; left M1: z=6.10, peak MNI coordinate x=-36, y=-18, z=48; left PMd: z=5.02, peak MNI coordinate x=-37, y=-6, z=55; right PMd: z=3.11 at x=37, y=06, z=55; **Figure 2A**). A further analysis performed on a 38-region parcellation using beamforming with orthogonalization ^30, 31^ confirmed that the beta values for both input and response suppression were strongest in the parcel containing premotor cortex (**Supplementary Fig 1**).

**Figure 2.**
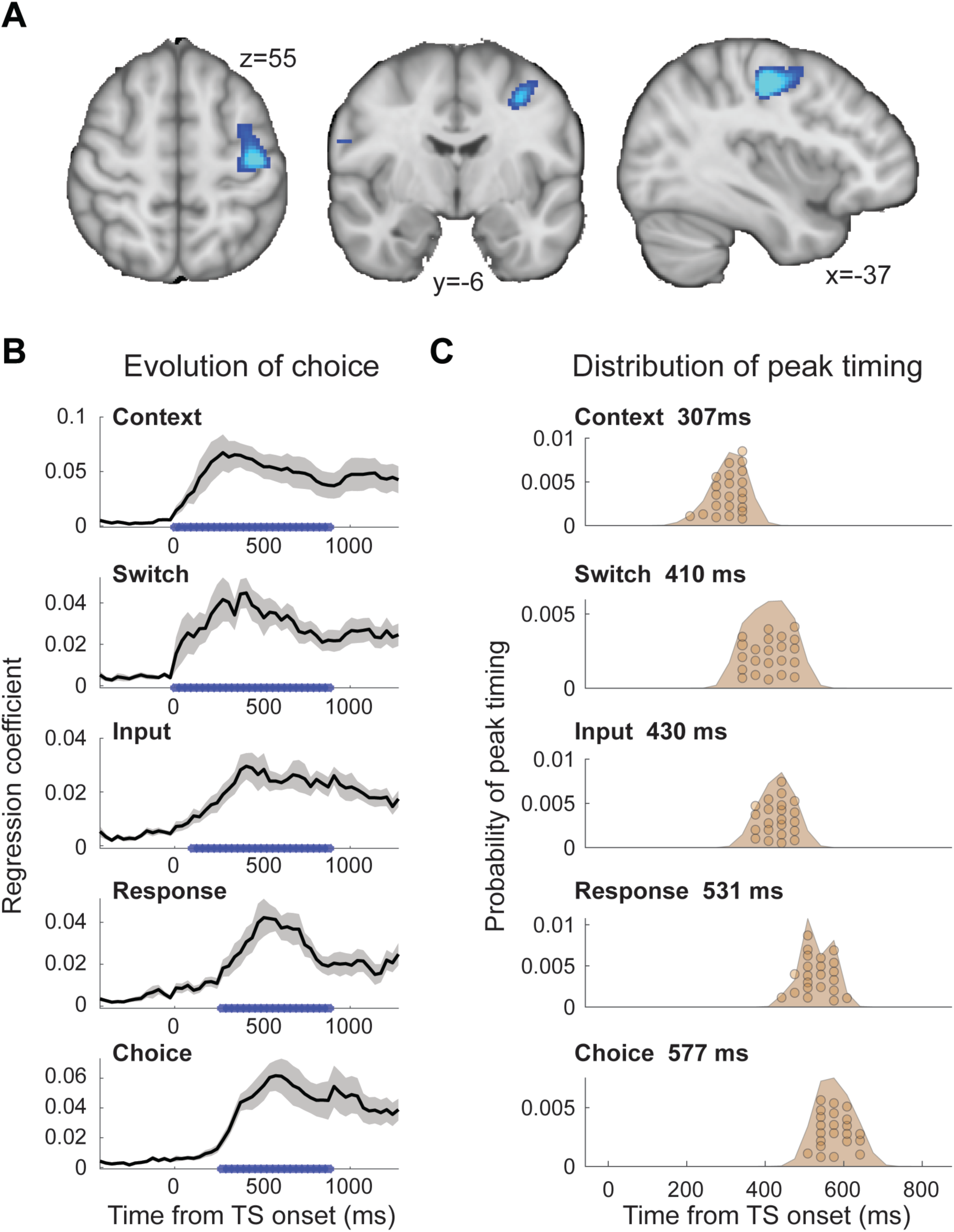
The evolution of choice in human premotor cortex. **A**, PMd was the *a priori* region of interest for this study. Indeed, beamforming for source localization identified a cluster involving M1 and left PMd (group contrast, whole-brain cluster-level FWE-corrected at *P*<0.05 after initial thresholding at *P*<0.001; PMd peak coordinate: x=-37, y=-6, z=55) from a contrast probing any input (relevant or irrelevant) or response adaptation. Importantly, this contrast was agnostic and orthogonal to any differences between the encoding of input and response adaptation, relevant and irrelevant input adaptation, or pre- vs post-training effects (including any timing differences, as beamforming was performed in the time-window [-250,750]ms). **B**, The time-resolved nature of MEG, combined with a selective suppression of different choice features, allowed us to track the evolution of the choice on a millisecond timescale. Beta coefficients from a regression performed on data from left premotor cortex (PMd) demonstrate an early encoding of context (motion or colour) and switch (attended dimension same or different from TS) from ∼8ms (sliding window centred on 8ms contains 150ms data from [-67,83]ms). The encoding of inputs emerged from around 108ms (whether or not the input feature, colour or motion, was repeated). Finally, the motor response (same finger used to respond to AS and TS or not) and choice direction (left or right) were encoded from 275ms. * indicates p<0.001; error bars denote SEM across participants; black line denotes group average. **C**, The distribution of individual peak times across the 22 participants directly reflects this evolution of the choice process. In particular, it shows significant differences in the encoding of input and response, consistent with premotor cortex transforming sensory inputs into a motor response.

To examine how the choice evolved over time in PMd, we extracted the timeseries from left PMd (x=-37, y=-6, z=55) and used a L2-regularized linear regression (ridge regression) containing regressors that described which task-variables were being suppressed on a trial-by-trial basis (see Methods). The regression was applied to each time-point around the presentation of the TS ([-500,1200]ms) using a sliding-window approach (window size: 150ms). **Figure 2B** demonstrates the temporal evolution of the choice for each of the regressors. The two contextual variables ‘context’ (motion or colour instruction flankers) and ‘switch’ (attended dimension same or different from AS) showed the earliest significant encoding and this response was sustained throughout the TS presentation (both first significant at 8ms). This is unsurprising as contextual information was available −150ms before the TS. Slightly later, starting from 108ms, the encoding of input emerged (probing whether or not the same input feature was already shown in the AS and thus suppressed). The regressor that was encoded latest in PMd was the motor response (probing whether the same finger was/was not used to respond to the AS and thus suppressed) and the choice direction (left or right; both starting from 275ms).

Statistical tests performed on the peaks of the beta values obtained for the five regressors showed a significant difference in encoding latency (one-way repeated-measures ANOVA: *F*(4, 84) = 128.71, *P* = 5.5154e-35, η^2^_p_ = 0.86; **Fig 2C**) and a pairwise post-hoc test between the two critical regressors, input and response repetition suppression, confirmed a significant timing difference with the encoding of choice input preceding the encoding of response direction (pairwise *t*-test on peak betas for input [429.5ms ± 35ms; mean ± s.t.d] and response [531.1ms ± 41.6ms]: *t*(21)=11.07, *P* = 3.1654e-9, after Bonferroni familywise error correction; *Hedges*’ *g* = 2.577; see Online Methods; **Fig 2C**). Thus, activity recorded using MEG repetition suppression in human premotor cortex can track the evolution of a choice from sensory input to motor response. This transition from input-to-choice encoding can also be plotted as a population trace, recapitulating the state space trajectories observed in previous NHP recordings ^4^ with input on one and response on the other axis (**Fig 3A**). This way of plotting the data mimics the dynamics of the neural population which normally relies on concatenating many neuron’s firing rates but which we obtained non-invasively. Thus, comparable insights into the population’s movement through state space during the decision process can be obtained.

**Figure 3.**
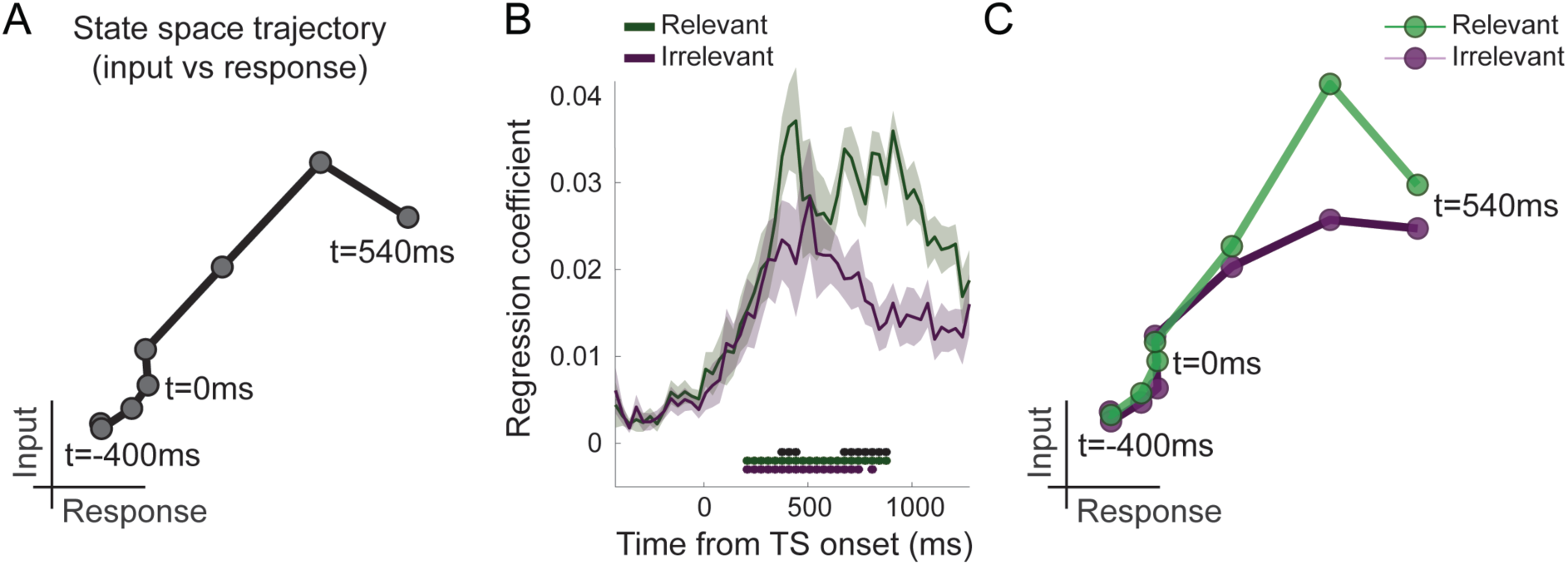
Population traces show filtering out of irrelevant inputs in PMd. **A**, By plotting the encoding of input on one and response on the other axis (both as in Fig 2B), we can derive a population trace, mimicking state space trajectories obtained from NHP recordings ^4^. **B**, Separation of input suppression into relevant and irrelevant inputs shows slightly diminished encoding of irrelevant inputs in PMd (*P*<0.001; black * show difference between relevant and irrelevant inputs: significant between 375-442ms and 675-875ms; green and purple * indicate significance separately for relevant and irrelevant inputs). **C**, This can also be seen in the state space population traces for relevant and irrelevant input. We observed partial but incomplete filtering of irrelevant inputs in PMd.

### Irrelevant inputs are encoded less relative to relevant inputs in human PMd

Having established that different components of the choice computation can be tracked in a temporally resolved manner using MEG, we next examined whether PMd processes inputs equally when they are relevant compared to when they are irrelevant for the choice at hand. Accounts of top-down attentional filtering would predict reduced encoding of inputs that are irrelevant for making a choice (e.g. colour when the choice is about motion). By contrast, strong evidence has been provided in recording from NHPs for the passing forward of both relevant and irrelevant inputs, with selection occurring at the output stage ^4^. Comparing the regression betas capturing the encoding of relevant and irrelevant inputs showed that both were encoded significantly from 208ms, but importantly relevant inputs were encoded more strongly compared to irrelevant inputs (*P*<0.001 between 375-442ms and 675-875ms; **Fig 3B**). This can also be appreciated in the state-space population traces for relevant and irrelevant input (**Fig 3C**). Thus, our data reconciles both accounts, with partial but incomplete filtering of irrelevant inputs in PMd. Feature selection did not occur solely at the motor output stage of the decision process, nor was irrelevant information entirely filtered out by this stage.

### Over-training does not abolish the filtering out of irrelevant inputs

We speculated that one potential cause for seeing attenuated irrelevant sensory input encoding in our human participants who spontaneously performed the task, but unattenuated irrelevant input encoding in recordings from NHPs, may be rooted in the fact that monkeys have trained to perform this task for many months and thousands of trials. To test this hypothesis, we trained our human participants for four weeks on over 20,000 trials of the task (see Methods; **Supplementary Figure 2A**). Unsurprisingly, this led to a speeding up of reaction times and higher performance scores (2 x 2 x 3 repeated-measures ANOVA with factors context [motion vs color], training [pre vs post] and coherence [70% (AS) vs. 25.6 vs. 12.8]; effect of training on log-RT: F(1,20)=9.23, p=6.46e-3, η^2^_p_ = 0.32, mean RT difference 29ms+/-10ms; effect of training on % correct: F(1,20)=36.39, p=6.75e-6, η^2^_p_ = 0.65, mean difference 4.76+/-0.8; **Fig 4A and Supplementary Figure 2B-E**). Upon completion of the training, we again recorded MEG data during performance of the task in the same way as done pre-training. As before, the unfolding of the choice computation was evident in signals recorded from PMd (**Fig 4C-D**). Crucially, however, when we repeated the analysis focussing on any differences between the encoding of relevant and irrelevant inputs, the difference was retained (**Fig 4E-G**). Even after having performed thousands of trials on the task, irrelevant inputs were therefore still filtered out from PMd activity. There was no interaction between input encoding and training (two-way repeated-measures ANOVA with factors input [relevant vs irrelevant] and training [pre vs post]: all *P*>0.05 for effect of input*training; Bonferroni correction for familywise error rate; a two-way repeated-measures Bayesian ANOVA provided further evidence in favour of the null hypothesis: the input*training interaction was not part of the winning model at any timepoint).

**Figure 4.**
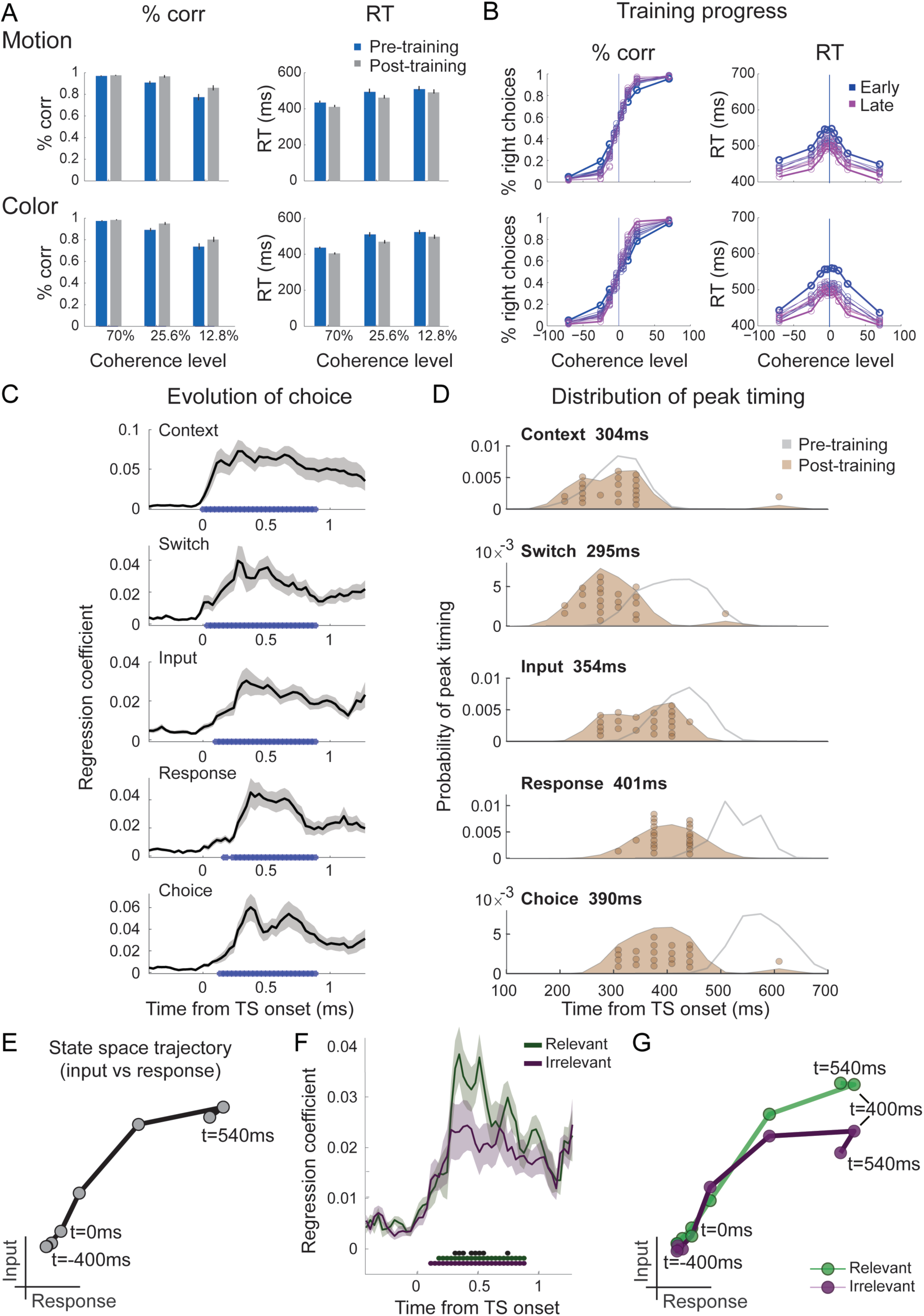
Filtering out of irrelevant inputs remains after overtraining on >20,000 trials. **A**, Over a period of four weeks, participants performed >1,000 trials of training on the random-dot-motion task on 5 or 6 days a week before returning for two further post-training MEG sessions. Comparison of pre- and post-training task performance showed significant RT speeding and performance improvements (% correct choices) in both color and motion contexts. **B**, Performance improvements are also clear in psychometric curves plotted across training sessions (blue=early, purple=late; see also **Suppl Fig 2D**). **C**, The evolution of a choice can be tracked in PMd post-training (as shown in Figure 2B for the pre-training sessions). **D**, However, peak timings of the majority of regressors occur earlier in the post-compared to the pre-training data suggesting a faster and possibly more efficient coding of the choice process. **E**, The population trace extracted from PMd shows a transformation from an encoding of inputs to an encoding of outputs, similar to Fig 3A for the data acquired pre-training. **F**, The difference between the encoding of relevant versus irrelevant inputs is preserved even after extensive training, thus suggesting some filtering out of irrelevant inputs. **E**, This can be visualized in two divergent population traces, separately showing relevant and irrelevant input adaptation.

Consistent with faster behavioural reaction times, we did observe changes in peak encoding latency of the different components of the choice process after completion of an extensive training regime (**Fig 4D**; two-way repeated-measures ANOVA with factors regressors [Context, Switch, Input repetition suppression, Response repetition suppression, Choice] and training [pre vs post]: effect of regressors F(4,84) = 39.35, *P*= 1.5824e-18, η^2^_p_ = 0.65]; effect of training [F(1,84) = 75.81, *P*= 2.0638e-08, η^2^_p_ = 0.78]; effect of regressors*training [F(4,84) = 35.38, P= 2.6436e-17, η^2^_p_ = 0.63]). Post-hoc tests confirmed that peak timings changed for four out of five regressors following the training, including the encoding of context switches, input adaptation, response adaptation and choice direction (pairwise *t*-test between pre vs post on the peak timings of context: *t*(21) = 0.18, *P* =1.00, *Hedges’ g* = .046; switch: *t*(21) = 5.92, *P* = 3.5641e-05, *Hedges’ g* = 1.98; input repetition suppression: *t*(21) = 5.91, *P* = 3.6398e-05, *Hedges’ g* = 1.57; response repetition suppression: *t*(21) = 12.15, *P* = 2.8914e-10, *Hedges’ g* = 3.08; choice: *t*(21) = 11.09, *P* = 1.5246e-09, *Hedges’ g* = 3.28; Bonferroni correction for familywise error rate; **Fig 4D**).

### Filtering out of irrelevant inputs is present across the brain

Finally, it is possible that our targeted focus on PMd as the ‘output’ stage of the decision process might have obscured the fact that other MEG sensors encoded irrelevant input information more strongly. To investigate this, we performed a multivariate decoding analysis on the data from all sensors. Consistent with earlier analyses, decoding across all sensors showed weaker decoding of irrelevant compared to relevant inputs (**Fig 5**). There was some significant decoding of irrelevant inputs (*P*<0.001 from 455ms to 708.3ms post-TS) but it was again less extended in time and weaker compared to the decoding of relevant inputs (*P*<0.001 from 201.7ms to 961.7ms post-TS; **Fig 5A**). There was again no difference between the data acquired pre- vs post-training (two-way repeated-measures ANOVA with factors input [relevant vs irrelevant] and training [pre vs post]: all *P*>0.5; minimum *P*=0.52, *F*(1,21) = 5.49, η^2^_p_ = 0.21, for effect of training; Bonferroni correction for familywise error rate; a two-way repeated-measures Bayesian ANOVA provided further evidence in favour of the null hypothesis: the input*training interaction was not part of the winning model at any timepoint) apart from differences in peak timing (**Fig 5B**; pairwise *t*-test between pre vs post on the peak timings of input repetition suppression: *t*(21) = 3.12, *P* = 0.026, *Hedges’ g* = 1.00; context: *t*(21) = 2.35, *P* = 0.14, *Hedges’ g* = 0.62; switch: *t*(21)= −0.60, *P* = 1.00, *Hedges’ g* = −0.19; response repetition suppression: *t*(21) = 1.59, *P* =0.63, *Hedges’ g* = 0.52; choice: *t*(21) = 2.30, *P* = 0.16, *Hedges’ g* = 0.67), as observed in PMd (**Fig 4D**). This further supports an interpretation of increased efficiency as a result of extensive over-training. Thus, while this analysis cannot rule out that there may be a brain region which encodes irrelevant inputs more strongly, we confirmed that even after extensive training and when considering the activity across the whole scalp, the processing of irrelevant inputs was attenuated compared to the processing of relevant inputs when making a choice.

**Figure 5.**
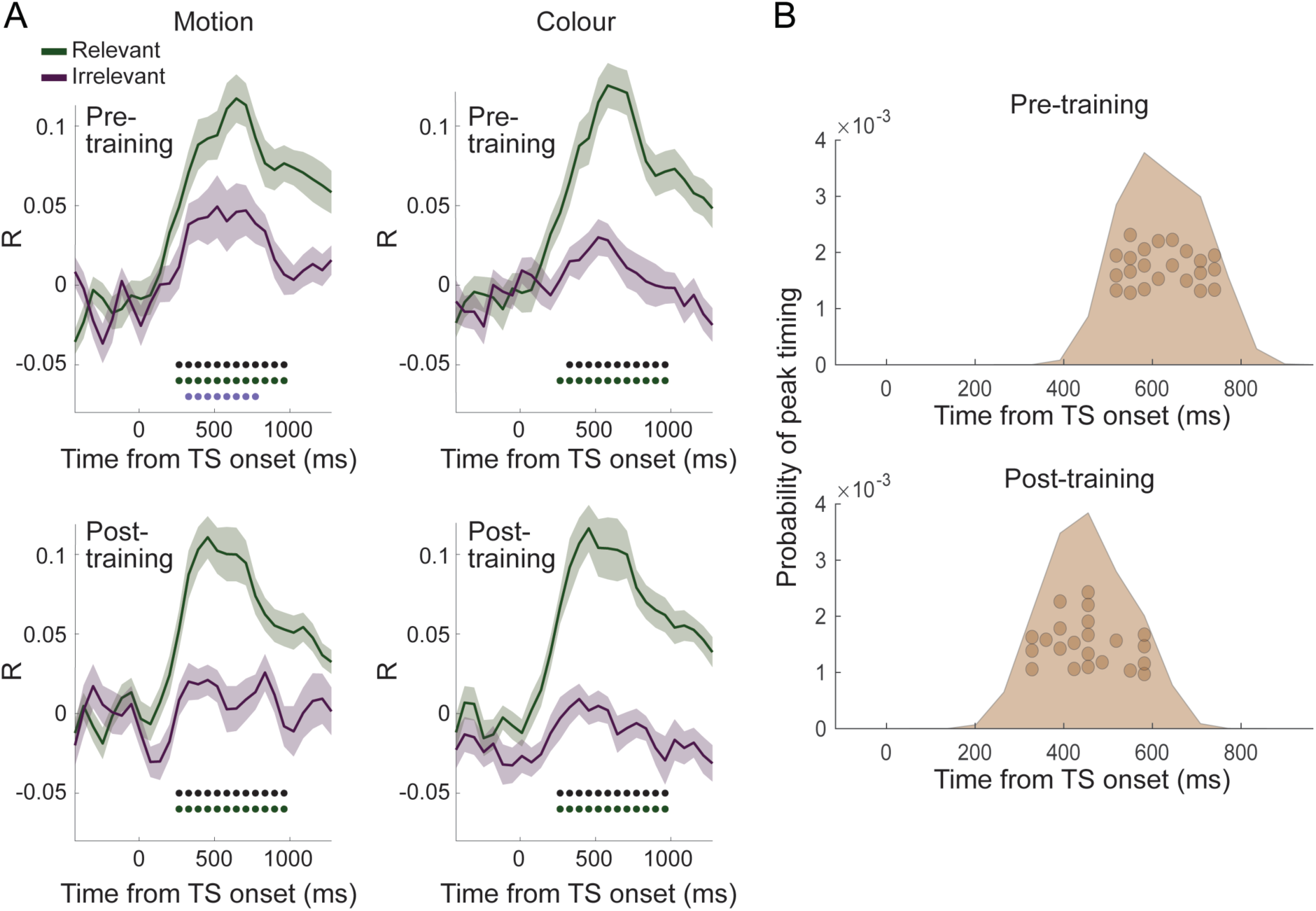
Filtering of irrelevant inputs is also present in whole scalp decoding. **A**, Decoding performed on all MEG sensors showed a significant difference in decoding accuracy between relevant and irrelevant inputs, suggesting an attenuation of information about irrelevant choice features. Thus, a filtering out of irrelevant information is apparent even when considering data from the whole brain, and it is not affected by extensive training on the task (compare top vs. bottom rows). As in Fig 3B, black * denote significance at *P*<0.001 for the difference between relevant and irrelevant input coding; green and purple * denote significance for relevant and irrelevant input decoding, respectively. **B**, Consistent with the encoding approach focussed on PMd, peak decoding times are faster following the training when considering data from all sensors.

## Discussion

Here we have shown that noninvasively recorded MEG activity can be used to track the state space trajectory of a choice process on a millisecond timescale. As decisions unfolded, MEG activity in premotor cortex transitioned from encoding sensory inputs to encoding the motor response. Watching these dynamics has previously only been possible by projecting high-dimensional neural population recordings onto a low-dimensional set of axes or ‘neural state space’. Here we took a different approach to measuring this state space, in which we selectively suppressed the encoding of either sensory inputs or response features in the average neural data recorded with MEG. This allowed us to selectively index neural activity along the different axes that define the neural state space. We found that in human premotor cortex, sensory inputs irrelevant to the current choice were encoded more weakly compared to relevant choice inputs. This partial filtering of irrelevant choice features was observed even after extensive over-training and when considering activity present across the brain.

Studying neural population responses, as opposed to the responses of individual neurons, has received increasing attention because it provides a window into larger-scale neural dynamics ^2^. It has provided crucial insights, for example, into the evolution of choice ^3–6, 32^, movement preparation and execution ^7–9^, and the mechanisms underlying working memory ^10^. The axes of the neural state space onto which the population response is projected are sometimes defined in a data-driven manner, as the population’s major axes of variation (e.g. ^7^). Alternatively, they may also be defined by projecting neural activity onto the relevant task variables, defined by each neuron’s regression weights to these variables. For example in ^4^, the three major axes of variation were the relevant and irrelevant sensory inputs and the choice direction. It is this latter form of population state space that we targeted in the current study.

The approach we use to derive this state space draws on the observation that the electrophysiological responses are suppressed when repeatedly exposed to features to which they are sensitive. In MEG data recorded noninvasively from human participants, each sensor’s signal is driven by the summed postsynaptic potentials of millions of neurons. This bulk signal can be manipulated by repeated exposure to a specific feature, causing suppression in the subset of neurons sensitive to this feature and thus reduce the average MEG signal that is measured (**Figure 1**). While this insight has been exploited for understanding a wide range of cognitive processes, with both fMRI ^33–42^ and MEG/EEG ^43–46^, here we extended this framework in one unusual way. As in previous work, using MEG allowed us to measure temporally resolved repetition suppression effects, thus giving insight into precisely when particular features (e.g. input versus choice direction) are processed. This has been exploited previously, for example, to characterize the temporal tuning of repetition suppression ^43^ or to temporally dissociate stimulus repetition from stimulus expectation effects ^46^. However, importantly, here we suppressed the signal to multiple different features within the same experiment (colour and motion - when relevant or irrelevant - and response direction). This allowed us to mimic projections onto multiple axes in state space, with each feature representing one axis.

The obtained human population traces closely resemble those from invasive recordings in NHPs (**Figs 3 and 4**) ^4^, showing a progression from an encoding of choice inputs to an encoding of the motor response. However, in ^4^ the influence of input returned to baseline before choice execution. This may be because monkeys were only allowed to indicate a response after a delay, once the motion stimulus had been turned off. By contrast, we allowed participants to respond at any time while the stimulus was presented, and thus the trial could end before input encoding returned to zero. Indeed, in our state space trajectories, input encoding started to return to baseline just before the time of response but did not fully return to zero by the time participants responded. Overall, we believe our approach provides an exciting new opportunity, for example, by allowing researchers to measure neural state space trajectories in tasks which might be difficult for NHPs to do (e.g., involving language), or in human disease.

Our data speak to an important controversy about the mechanisms underlying feature-based, or context-dependent, choice selection. A substantial body of evidence has shown that selection of relevant sensory features occurs through top-down modulations from prefrontal and parietal regions onto early sensory regions ^13, 14, 16–18, 20–27, 47^. By contrast, the study by ^4^ found that irrelevant sensory features were not filtered out but passed forward to the output stage see also ^28^. The authors provided a compelling neural network model that solved feature selection and evidence integration within a single recurrent network. Because the monkeys in ^4^ were extensively over-trained at performing the task, we hypothesized that top-down influences might be more important when someone is naïve at doing the task. Indeed, in a task with only relevant features, training changes neuronal responses responsible for interpreting sensory evidence, but not those processing the sensory evidence itself ^48, 49^. We therefore compared participants before and after >20,000 trials of training on the task. To our surprise, the only difference identified between naïve and extensively over-trained participants were shifts in peak timing (**Figs 4 and 5**). We did not observe any changes in feature selection. In other words, even following the training, irrelevant features were present but weaker than relevant features in premotor cortex and across the brain. Our data thus suggests partial but incomplete filtering of irrelevant inputs in PMd. This is broadly in line with the effects present in participant’s behaviour, where we observed a weak but significant influence of the irrelevant feature (**Suppl Fig 2D**). Consistent with predictive coding principles and theories of top-down control, the processing of irrelevant inputs was diminished, but partly consistent with ^4^, they were not completely filtered out. Thus, it seems plausible that some but not all of the feature-selection happens at the output stage, in PMd. Overall, the strength of top-down control, or the extent to which task-irrelevant information was filtered out, seemed unaffected by the amount of prior experience on the task.

This leaves outstanding the question as to why equally strong encoding of relevant and irrelevant features were observed in NHP state space trajectories (Mante et al., 2013), but not in our data. It has been proposed that the site of feature selection may depend on the level of detail afforded by the prediction ^50, 51^. One difference between tasks was that the relevant dimension changed from trial to trial in our experiment but was blocked in ^4^. There is evidence that FEF shows long-term selection history effects ^52, 53^ which may be promoted by blocking trials. However, ^28^ do not mention any blocking of trials and report results consistent with ^4^, making this an unlikely possibility. Recent work has shown attenuation of expected stimulus features when they are attended ^54^. However, attenuation of expected information is usually thought to help filter out predictable objects ^16, 55^, for example via representations of pre-stimulus sensory templates ^56^. Indeed, generally, processing is biased in favour of behaviourally relevant input for review, see e.g.,^14, 57^. Ultimately, the discrepancy between different findings remains to be addressed and highlights a general need for a better understanding of decision-making in environments that require dynamic changes ^58–60^.

Overall, our results reinforce the importance of inter-species translational research, whereby tasks and techniques are used across species e.g. by using comparable analysis pipelines ^61, 62^, by obtaining direct recordings from humans when possible (e.g. neurosurgical patients: ^63–67^), or by recording whole-brain fMRI in NHP species ^68, 69^. Our work also emphasizes the importance of developing more mechanistic approaches in human neuroscience, and it shows that the generalizability from NHPs to humans can and should be tested but not assumed ^70^.

## Methods

### Participants

Twenty-five participants (10 male, 15 female, age range 19-32, mean age 25 ± 0.68) with no history of neurological or psychiatric disorder, with normal or corrected-to-normal vision and who fulfilled screening criteria for undergoing MRI and MEG scanning took part in this study. One participant dropped out before completing the experiment and two participants’ data sets were too noisy even after rigorous data clean-up. The final sample thus included twenty-two participants (10 male, 12 female, age range 19-32, mean age 25 ± 0.74). There was a problem with processing the MEG data from two sessions (session 2 and session 4 in two different individuals) so only data from three instead of four MEG sessions were included for two participants. The study was approved by the University College London (UCL) Research Ethics Committee (reference 1825/005) and all participants gave written informed consent.

### Experimental procedure

Participants agreed to take part in an initial screening session which ensured that they were safe to undergo MRI and MEG scanning, and that they were able to do the task to a basic standard (e.g., they were not color blind). No participants were excluded after the screening session. Following the screening, they then took part in a short structural MRI scan and four MEG sessions, two at the beginning and two at the end of the study, spaced four weeks apart. They also agreed to complete an hour of training (including short breaks: 1200 trials split into 12 blocks) in the laboratory or on their own computers on five or six days per week for a total of twenty-two training sessions spread across four weeks (**Supplementary Fig 2A**). Participants who performed their training at home (9 out of 22) agreed to pass on the data to the experimenter on the same day to enable monitoring of progress and to ensure daily completion, and they agreed to perform the task in a quiet environment without interruptions. Personal laptop screens were color-calibrated to ensure matched stimulus appearance. The four MEG sessions were identical and lasted ∼1.5h (1024 trials). Participants were reimbursed £250 for their time. Half of the money (£125) was paid out in smaller chunks after each MEG session and each week of behavioral training; the other half was paid upon completion of the entire experiment to discourage drop-out given the time-intensive nature of this study.

### Experimental task

The task was adapted from ^4^ and contained the same random-dot-motion stimuli (RDM) that in addition to a dominant motion direction also contained color information, here varying from predominantly green via grey (neutral) to predominantly red. Each trial contained two sequentially presented colored RDM stimuli, the adapting stimulus (**AS**) and the test stimulus (**TS**). Two separate instruction cues, presented shortly before the AS and TS, signaled whether participants had to judge the direction of motion or the color dominance of the AS and TS, respectively. This determined the relevant input dimension to focus on. More precisely, within a given trial, the order of presentation was as follows (**Figure 1A**): (1) an instruction cue presented for 150ms showed a green and red dot to signal that color was relevant or a left and right arrow to signal that motion was the relevant dimension to attend to for the AS; (2) The AS was presented for 500ms, either with random motion and 70% color dominance for green or red, or with non-dominant color (grey-shades) and 70% motion coherence to the left or right; (3) a fixation cross was shown for the 300ms inter-stimulus-interval (ISI); (4) a second instruction cue signaled the relevant dimension for the TS (150ms); (5) the TS was presented for 500ms; (5) after another 500ms of fixation, (6) feedback was presented for 300ms (‘green tick’ or ‘red cross’). Participants had to respond to both AS and TS using a button press with their right-hand index (left) or middle (right) finger. Because the response to the AS was trivial (dominance level: 70%; accuracy 95% +/- 2% during screening), the feedback at the end of the trial related to their response to the TS. TS color and motion dominance were modulated according to two difficulty levels during the MEG sessions: 12.8% or 25.6%. During the training, we also included two other difficulty levels corresponding to 3.2 and 6.4%.

During the screening session, the four MEG sessions and the last three days of training, this was the precise task used. During the screening session, participants performed six blocks of 128 trials (n=768 trials) of the task. During MEG sessions, they performed 8 blocks of 128 trials and thus a total of 1024 trials each, allowing a total of 2048 trials from each participant to enter the pre- vs post-training MEG analyses. During the first seven days of behavioural testing following the first two MEG scans, a simpler version of the task was used. Participants were only given one stimulus at a time (coherences: 3.2, 6.4, 12.8, 25.6 and 70%) and it only contained either color or motion (‘1-dimensional’ stimuli, 1D; **Supplementary Fig 2A**). There was feedback after every stimulus and even though it was easy to know which feature to attend to (when all dots were grey, it was motion, when they were colored and static, it was colour), the instruction cue was presented 150ms prior to stimulus onset. Participants performed 12 blocks of 100 trials (n=1200) per day, totaling to 8400 trials across the seven days. Following the 1D task, participants moved on to individual 2D stimuli that simultaneously included color and motion and performed this 2D task for 12 days (coherences were identical to the 1D task). Again participants performed 12×100=1200 trials per day, totaling to 14,400 trials of this 2D version of the task. Finally, the last three days of training were done on the task described above containing two stimuli presented in quick succession (AS=70% coherence and TS=12.8% or 25.6% coherence), and which was identical to the one used during the MEG scans (8 blocks of 128 trials per day and thus 3 x 1024 = 3072 trials in total). Thus, all participants were expected to complete a total of 7+12+3 = 22 training sessions. They were told not to take more than one day off in a row but due to sickness, some sessions are missing in some participants (mean number of completed sessions: 21.2). Overall, by the time they came for their third and fourth MEG session, participants were expected to have completed 768 (screening) + 2048 (2 MEG) + 8400 (7 days 1D stimuli) + 14400 (12 days 2D stimuli) + 3072 (3 days full task) = 28,688 trials. Everyone performed the screening, the four MEG sessions, and all seven 1D sessions. Of the twelve 2D sessions, participants completed between 5-12 (mean: 10.7) and of the final three full task sessions, they completed between 1-3 (mean: 2.5) before coming back for the two post-training MEG sessions. In total, everyone completed >20,000 trials before the final MEG sessions and on average 26,594 trials (minimum: 20,288, maximum: 28,688).

### Repetition suppression procedure and trial types

All analyses focus on the time of the TS. Importantly, however, the purpose of the AS was to selectively manipulate neurons responding to particular input and response features. For instance, presenting a green AS followed by a predominantly green TS meant that at the time of TS, any MEG sensors influenced by neurons responding to green color, or by neurons responding to leftward hand motor responses should show suppressed responses compared to a situation where a red AS was followed by the same predominantly green TS. In a similar way we could selectively adapt to green or red color inputs, right or left motion inputs and middle/index-finger hand motor responses, and we could do so for when a given input was relevant or irrelevant. For example, a green AS followed by a left-motion TS that was predominantly green but while attending to motion, suppressed to green color when it was irrelevant at the time of TS. Finally, response adaptation could be obtained e.g. by showing a red AS (leading to a right and thus middle finger response) followed by a right-moving TS (also leading to a response with the middle finger). The full table of conditions can be seen in Table 1. In total there were 64 conditions: 4 AS x 2 TS contexts (color/motion) x 2 TS directions (right/left) x 2 TS colors (green/red) x 2 TS coherence levels (12.8 and 25.6%). Trials of all types were interleaved and shown in a random order.

### Stimulus generation

Custom-written MATLAB (The MathWorks, Inc., Natick, Massachusetts, US) code was used to produce a randomized stimulus order for each session and subject, with balanced trial numbers for each of the 64 conditions. For each random-dot stimulus, a new random dot placement was generated, and given the appropriate level of coherence and motion. The RDM stimuli were coded using three interleaved streams of stimuli, one per screen refresh rate (17ms), with the following parameters: speed of dots: 4 degrees/second; temporal displacement 50ms (3 screen refreshs); spatial displacement: 0.2 degrees/second; unmasked area: 10×10 degrees; dot diameter 0.3 degrees; and number of dots: 100. The stimulus presentation was programmed in MATLAB and performed using the Psychophysics Toolbox ^71^.

### Behavioural analysis

We recorded choice (left or right button press) and reaction time (RT) to both AS and TS in each trial. To examine training improvements, average RT and % correct from the two pre-training MEG sessions were compared with those obtained in the two post-training MEG sessions (which used the same stimuli/schedule; black in **Supplementary Fig 2A**; see also **Supplementary Fig 2B-E**). We used an ANOVA with factors coherence (70%=AS, 25.6%=easy TS and 12.8%=hard TS), context (color or motion), and training (pre vs post) to assess statistical significance (**Figure 4A**).

### MEG and MRI data acquisition

MEG data was recorded continuously at a sampling rate of 600 samples per second using a whole-head 275-channel axial gradiometer system (CTF Omega, VSM MedTech). Participants were seated upright in the scanner and their head location was monitored using three fiducial locations (nasion, left and right pre-auricular points). The distance to the screen was measured to adjust the size of the stimuli and the lights were turned off. Eye movements were recorded (EyeLink software) which required a brief calibration and validation procedure. During each MEG session, participants then performed eight blocks of seven minutes of the task, with short breaks in-between. Responses were indicated using a keypad with their right-hand index and middle finger. All four MEG sessions (two pre-training and two post-training) were identical in terms of the difficulty, trial structure and procedure. One of the MEG sessions was followed by a short MRI session during which a structural T1-weighted MPRAGE scan was acquired on a 3T Magnetom TIM Trio scanner (Siemens, Healthcare, Erlangen, Germany) with 176 slices; slice thickness=1 mm; TR=7.92 ms; TE=2.48 ms; voxel size=1×1×1 mm.

### MEG data preprocessing

MEG data were preprocessed using SPM12 (http://www.fil.ion.ucl.ac.uk/spm/) and custom-written MATLAB code. Data were converted to SPM12 format, a notch filter was applied, and eyeblinks were removed based on the electro-oculogram channel using a regression procedure based on the PCA of the average eye blink as previously explained in ^72^. The data was downsampled to 300Hz, epoched at [-1500,2000] around TS onset, and baseline corrected between [−1500 −1100] (thus using a pre-AS baseline). Where necessary, timings were corrected for one frame (1/60s) between trigger and image refresh, which was based on timings recorded with an in-scanner photodiode. Trials containing artefacts were rejected visually using Fieldtrip’s spm_eeg_ft_artefact_visual. Prior to source localization, data were low-pass filtered at 40Hz and the blocks from each session were merged.

### MEG source reconstruction

Source reconstruction was performed in SPM12. The structural scans were segmented and normalized to the MNI template. A subject-specific mesh was created using the inverse normalization and the three recorded scalp locations were registered to the head model mesh. A forward headmodel was estimated for each session and subject (EEG BEM, single shell). A linearly constrained minimum variance (LCMV) beamformer was applied in the window [-250 750ms] around TS to estimate whole-brain power images on a grid of 5mm and for source data (virtual timecourse) extraction, using PCA dimensionality reduction to regularize the data covariance estimation ^73^. Although beamforming has proved powerful at reconstructing source signals in electromagnetic imaging, it can be limited in the presence of highly correlated source signals, such as those that can occur bilaterally across hemispheres. To overcome this, a bilateral implementation of the LCMV beamformer was employed, in which the beamformer spatial filtering weights for each dipole were estimated together with the dipole’s contralateral counterpart ^74^. Beamformed power images from the two pre-training sessions and the two post-training sessions were smoothed, log-transformed and averaged, respectively.

### Region of interest

The *a priori* region of interest for this study was dorsal premotor cortex (PMd). Two analyses performed on our data justified the choice of PMd. First, we ran a broad inclusive beamforming contrast on any adaptation (whether color or motion or response, relevant or irrelevant), averaged across pre- and post-training MEG sessions to avoid bias in subsequent analyses, and identified PMd within the peak cluster (p < .05, family-wise error (FWE) cluster corrected across the whole brain after initial thresholding at p<.001). Left PMd (x=-37, y=-6, z=55) was then used for extraction of time courses and further analyses on PMd source data (all subsequent statistical tests were orthogonal to ROI selection). Second, we used an established parcellation that included orthogonalization (to remove spatial leakage between parcels) to extract source data from 38 parcels obtained from an ICA decomposition on resting-state functional magnetic resonance imaging data from the Human Connectome Project ^30, 31^ and confirmed that the strongest task-related effects were present in the parcel that contained left PMd (**Supplementary Figure 1**).

### Linear regression on MEG source data in PMd

We fitted an L2-regularized linear regression (ridge regression) to the raw source data extracted from PMd, which contained the following six regressors capturing task events and repetition suppression effects:

1. Context [1/0; Motion/Colour]
2. Switch instruction [1/0; Switch/No-switch]
3. Relevant input adaptation [1/0; Motion or Colour relevant input adaptation/No relevant input adaptation]
4. Irrelevant input adaptation [1/0; Motion or Colour irrelevant input adaptation/No irrelevant input adaptation]
5. Response adaptation [1/0; Response adaptation/No adaptation]
6. Choice [1/0; Right/Left]

The regression was applied to each time-point around the presentation of the TS ([-500, 1350]ms) for each subject. To increase sensitivity, we used a sliding-window approach by averaging time-points within 150 ms around the timepoint and used a step-size of 33.3 ms. For each time-point, we sub-sampled 90% of trials from all correct trials. We then fitted ridge regression (MATLAB’s function fitrlinear) to this sub-sample for obtaining regression coefficients. Because ridge regression has a hyper-parameter λ (the regularization coefficient), we tuned λ from {10^-5^, 10^-3^, 10^-1^, 10^1^, 10^3^, 10^5^} using 3-fold cross validation for each sub-sample. Specifically, we used the hyper-parameter λ which performed the best in the 3-fold cross validation for estimating regression coefficients for the subset. We averaged regression coefficients across the ten sub-sets’ results for obtaining the estimated regression coefficients used for the following statistical analyses. To test whether regression coefficients were significant, we generated a null distribution by shuffling the trials within subject. We generated n = 1,000 permutations such that correspondence between trials and regressors were shuffled. We estimated regression coefficients using exactly the same procedure as above using these shuffled data.

We used a conservative time window to correct for multiple comparisons across time. Because the context cues came on at −150ms and responses were on average at around 500ms, we chose a window of nearly 1s duration from [-133.3, 950] ms around the TS. At our sampling resolution, this window contained 29 datapoints, and we corrected all statistical tests on these data across these 29 data samples. This correction was used to establish significance of individual effects (e.g. response adaptation) or differences between two effects (e.g. relevant versus irrelevant input adaptation). Note that, with alpha set at 0.05, and 29 time-points, the *p*-value required for significance would be 0.05/29 = 0.0017 after Bonferroni familywise error rate correction. Therefore we used a threshold of *P*<0.001 in the main figures.

We used the absolute values of the regression coefficients for statistical analyses and figures because source-localized MEG data have an arbitrary sign as a consequence of the ambiguity of the source polarity. As beamforming is done for each session separately, the sign of the reconstructed dipoles risks being inconsistent across subjects and sessions. We aligned the signs within subject for the two pre-training and the two post-training sessions separately by calculating Pearson’s correlation coefficient between average ERPs ([-200, 1500]) for session 1 and 2, and 3 and 4. In case of a negative correlation, we flipped one session’s signals before estimating regression coefficients. This ensured that pre- and post-training sessions each used sign-aligned data in a given participant.

To establish whether peak timings between input and response adaptation differed, the peak time for the parameter estimates were established in each participant for both regressors using an out-of-sample procedure. The average peak time across all participants except the left-out participant was determined and the left-out participant’s peak was taken as the highest regression coefficient in a window of size [-66.7, 66.7] around that group peak. We confirmed that using a wider window of size [-133.6, 133.6] ms did not change our conclusion. This procedure was repeated for all participants and all regressors and peak times were subjected to a one-way repeated-measures ANOVA. We conducted post-hoc pair-wise *t*-test between the peak timings of betas for input and response adaptation. For Fig 2C, a probability distribution of the peak time for each regressor was estimated using the fitdist function in MATLAB for visualization purposes.

Population traces were generated by simply plotting the regression coefficient obtained for input and response adaptation against each other within the same plot.

To investigate training effects on relevant versus irrelevant input encoding, we used a two-way ANOVA on the estimated regression coefficients with factors training (pre vs post) and input adaptation (relevant vs irrelevant) for each time-point in the window [-133.3, 950] ms. Bayesian ANOVAs were performed in JASP (JASP Team (2018), https://jasp-stats.org) and were JZS Bayes factor ANOVAs ^75, 76^ with default prior scales and enabled measuring the likelihood of the null hypothesis. Across time points, the largest P(M|data) for the model including the input*training interaction in PMd was P(M|data)=0.166. The winning models were either the Null model (17 time points) or a model with only a factor of input (relevant vs irrelevant; 9 time points) and their P(M|data) was >.4 across time (mean 0.52).

### Decoding from whole-brain MEG scalp data

Finally, to rule out that we were overlooking potential irrelevant adaptation effects by focusing solely on PMd, we repeated a similar analysis on the whole-brain MEG scalp data. A decoder was constructed separately for each time-point around the presentation of the TS ([-506.7,1416.7] ms) for each subject and each session. We used signals from 38 parcels (described above) for predicting current sensory evidence which takes one of four values from [–2, −1, +1, +2] indicating the level of positive or negative sensory evidence. For each time-point and sensor, choice information (right or left) was regressed out of the signal. The sign of the motion and color coherence is defined such that positive coherence values correspond to evidence pointing towards a right choice, and negative values correspond to evidence pointing towards a left choice. To increase sensitivity, we used a sliding-window approach by averaging time-points within 150 ms around the timepoint and used a step-size of 63.3 ms. For each context (motion and colour), we constructed a decoder for relevant input (e.g. motion input in motion context) and irrelevant input (e.g. motion input in colour context) separately. We used ridge regression and decoding performance was evaluated using a nested cross validation procedure as follows. We first split all correct trials into 10 sets of trials (10-fold outer-CV). We then split whole-trials except 1 held-out set into 3 sets of trials (3-fold inner-CV). We tuned the hyper-parameter λ from {10^-5^, 10^-3^, 10^-1^, 10^1^, 10^3^, 10^5^} in this 3-fold inner-CV. Finally, we fitted the model with the best-performing λ from the inner-CV to 3 sets of trials and obtained the prediction for the 1 held-out set. This procedure was repeated 10 times. Here, we chose a window of nearly 1s duration from [-190, 1036.7] ms around the TS for statistical testing. At our sampling resolution, this window contained 18 time-points.

Again, Bayesian ANOVAs were performed in JASP. Across time points, the largest P(M|data) for the model including the input*training interaction in the scalp data was P(M|data)=0.273. The winning models were either the Null model (2 time points), a model with only a factor of input (relevant vs irrelevant; 10 time points) or one with timing and input factors but not their interaction (4 time points). Their P(M|data) was >.3 across time (mean 0.56).

## Author Contributions

TEJB and MCKF designed the task, MCKF acquired the data, MCKF and YT analyzed the data, LH, MWW and TEJB advised on analyses and all authors wrote the paper.

## Acknowledgements

We would like to thank Gareth Barnes and Vladimir Litvak for advice on initial data analyses, the whole support team at the FIL for help with data acquisition, and MaryAnn Noonan, Nick Myers and Lev Tankelevich for helpful discussions on the manuscript.

## Data and code availability statement

The data and code supporting the findings of this study are available from the corresponding author upon reasonable request.

---

**Supplementary Figure 1.**
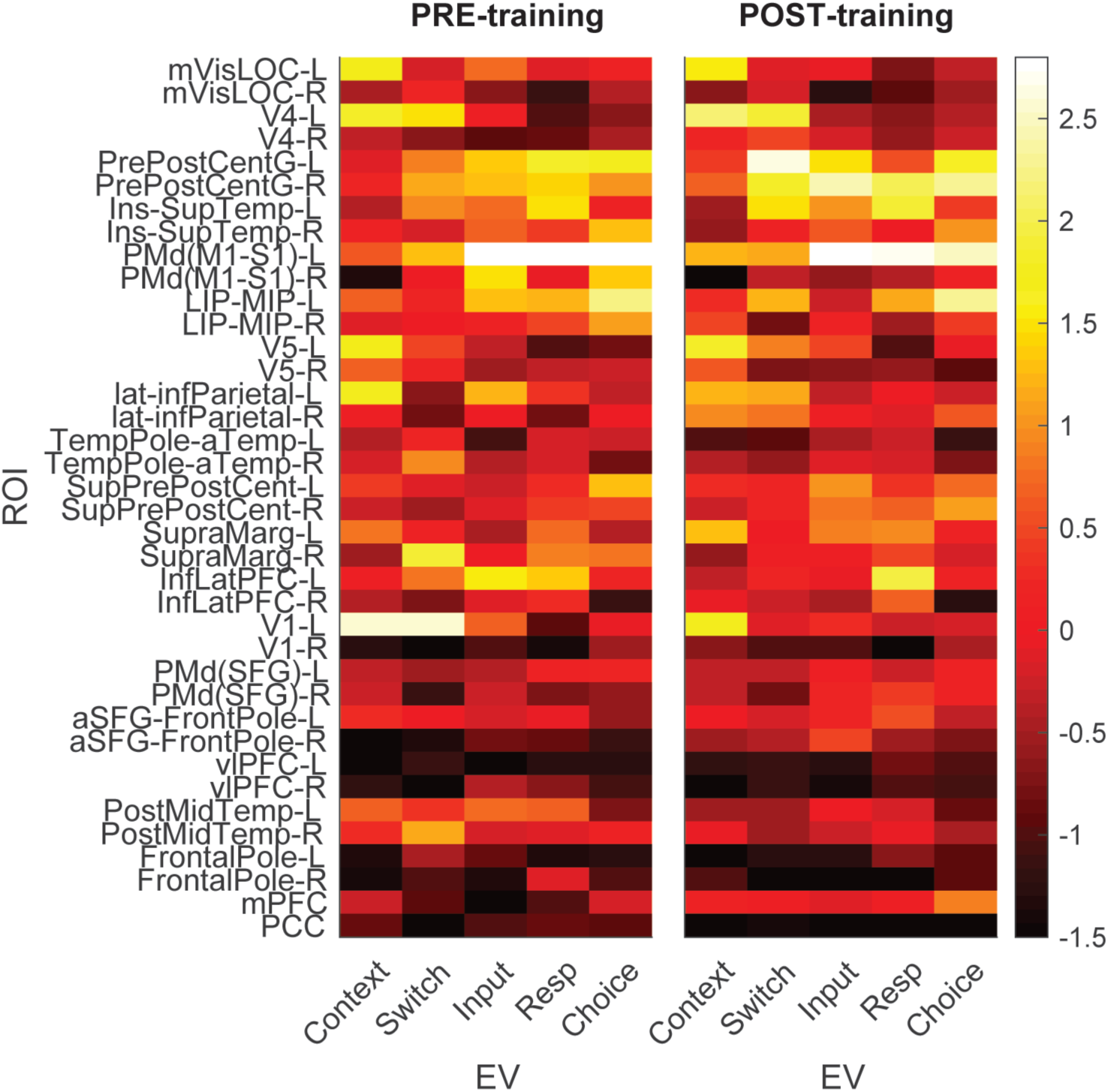
Premotor cortex encodes choice inputs and outputs. PMd was the *a priori* region of interest for this study. As shown in Fig 2A, beamforming for source localization identified a cluster involving left PMd from a contrast probing any input or response adaptation across both pre- and post-training sessions. Here we show an additional parcellation into 38 regions using beamforming which confirmed that the beta values for both input and response suppression were strongest in the parcel containing premotor cortex (PMd(M1-S1)-L).

**Supplementary Figure 2.**
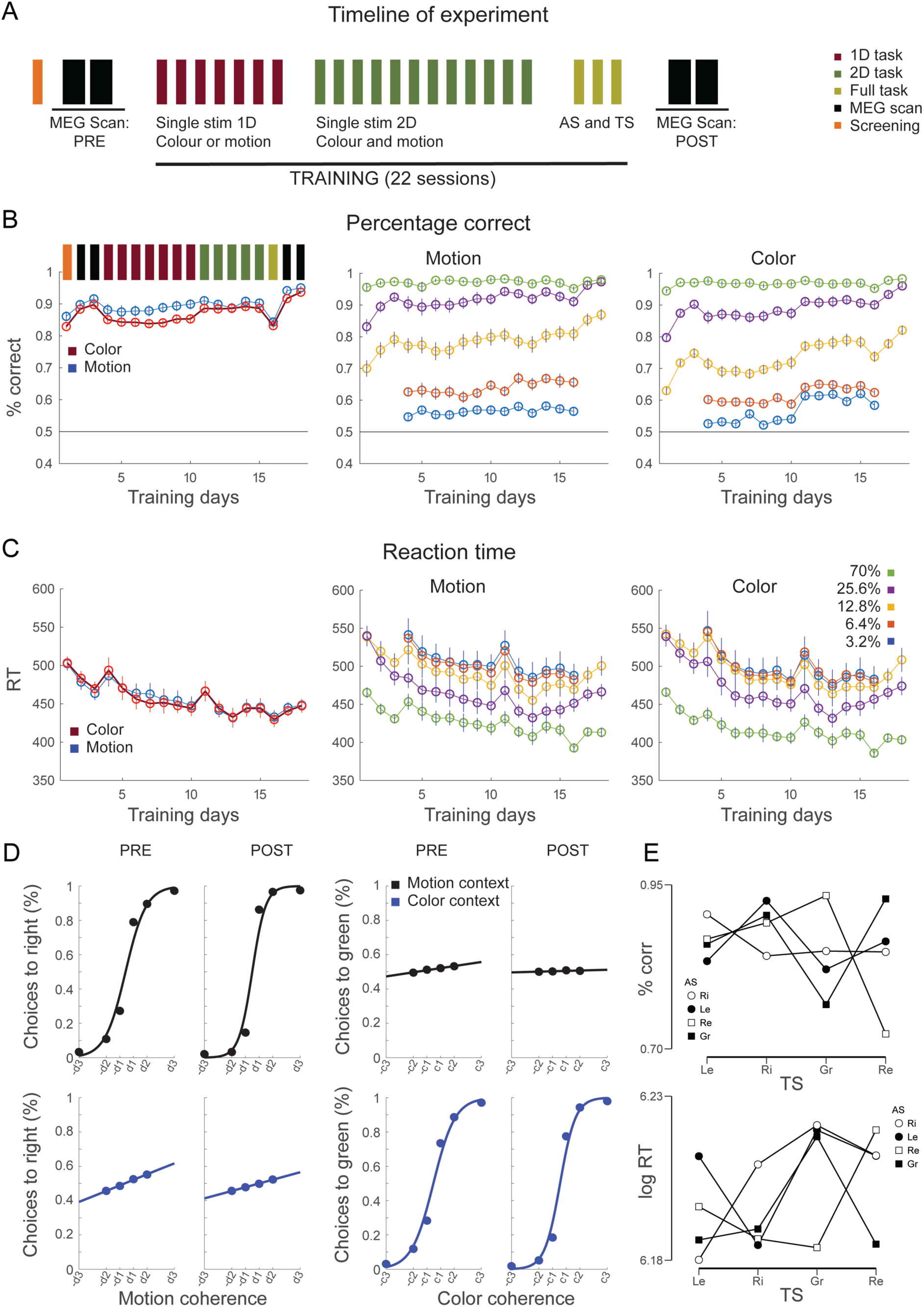
Timeline, performance progression during training sessions and behavioral effects of relevant and irrelevant input and AS. **A**, The timeline of the experiment is illustrated. The manuscript focusses on the neural data acquired from the two PRE-training and the two POST-training MEG scans (black). The training performed in-between was performed in three stages, starting with RDM stimuli containing only one dimension (color or motion: 1D), progressing onto RDM stimuli containing both dimensions (color and motion: 2D), followed by three sessions that contained 2D stimuli pairing the AS and the TS. Not all participants completed all 7 1D + 12 2D + 3 (AS/TS) = 22 training sessions. Everyone performed the screening, the four MEG sessions, and all seven 1D sessions. Of the twelve 2D sessions, participants completed between 5-12 (mean: 10.7) and of the final three full task sessions, they completed between 1-3 (mean: 2.5) before coming back for the two post-training MEG sessions. In total, everyone completed >20,000 trials before the final MEG sessions and on average 26,594 trials (minimum: 20,288, maximum: 28,688). **B**, Choice performance (% correct) is shown for the sessions completed by all participants, split by motion and colour (left), by coherence level for motion (middle) and for color (right). **C**, The same progression as in C is shown for reaction times (RT). **D**, Psychometric curves show that a strong influence of the relevant feature (color in a color trial and motion in a motion trial) and a weak influence of the irrelevant feature (color in a motion trial and motion in a color trial) were present both pre and post training, consistent with data in non-human primates (Mante et al., Nature, 2013) and consistent with the presence of the corresponding signals in the neural data recorded from PMd. **E**, The AS elicited contrast effects in the performance at the time of the TS. *Top*: accuracy was improved when TS features were contrasting with AS features, for example when a green AS was followed by a red TS (Ri=Right Motion, Le=Left Motion, Re=Red Color, Gr=Green color). When AS and TS shared features, accuracy was impaired, for example for a left motion AS followed by a left motion TS. *Bottom*: Similar effects were observed for the log-reaction time to the TS. Repeated motion or color slowed RTs, whereas contrasting motion or colour sped up RTs.

**Supplementary Table 1.**
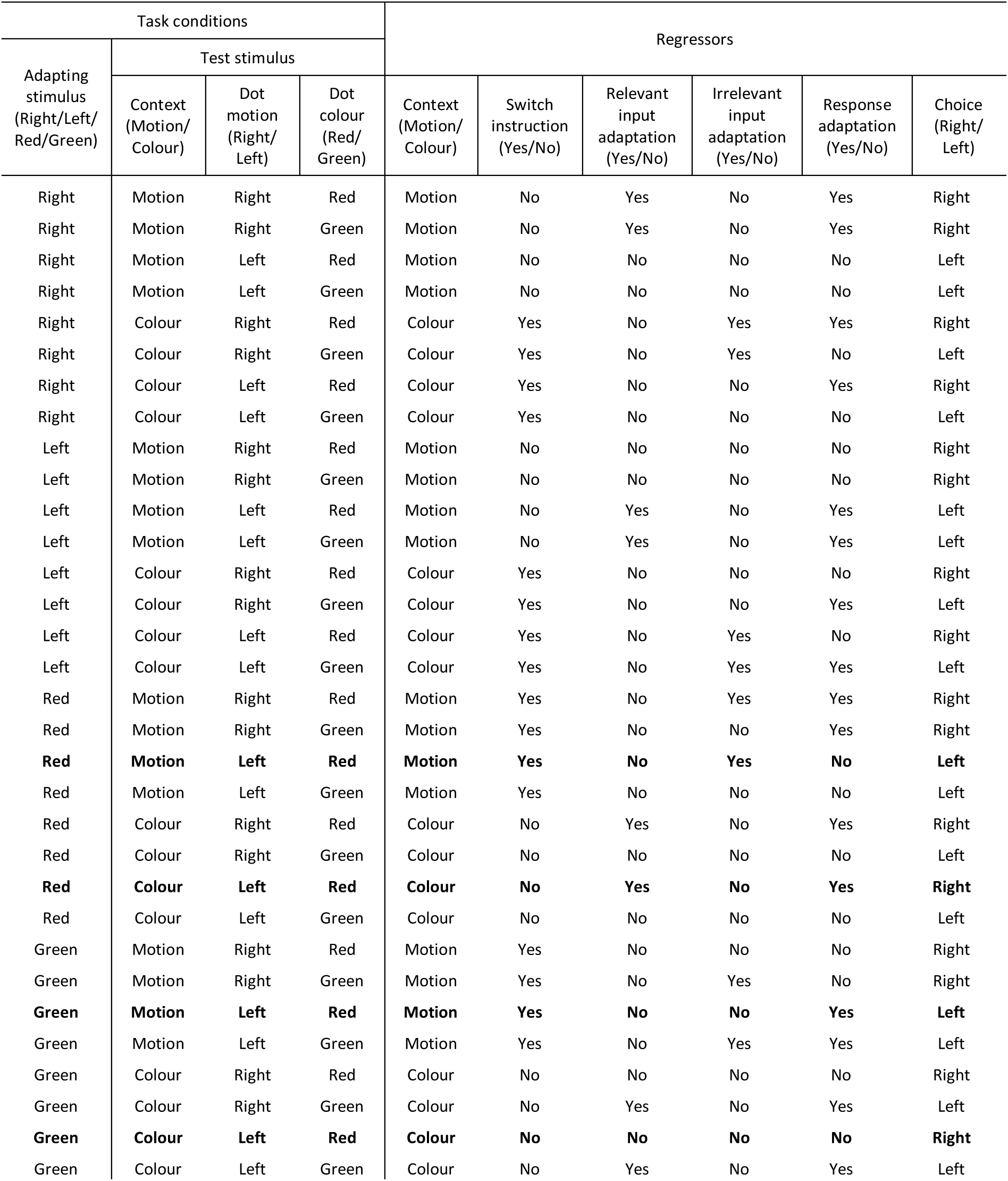
Task conditions and the corresponding regressors. The list of task conditions and corresponding regressors of the experiment are shown. The four bold lines are illustrated as examples in Figure 1B.

